# Arousal impacts distributed hubs modulating the integration of brain functional connectivity

**DOI:** 10.1101/2021.07.12.452041

**Authors:** Kangjoo Lee, Corey Horien, David O’Connor, Bronwen Garand-Sheridan, Fuyuze Tokoglu, Dustin Scheinost, Evelyn M.R. Lake, R. Todd Constable

## Abstract

Even when subjects are at rest, it is thought that brain activity is organized into distinct brain states during which reproducible patterns are observable. Yet, it is unclear how to define or distinguish different brain states. A potential source of brain state variation is arousal, which may play a role in modulating functional interactions between brain regions. Here, we use simultaneous resting state functional magnetic resonance imaging and pupillometry to study the impact of arousal levels indexed by pupil area on the integration of large-scale brain networks. We employ a novel sparse dictionary learning-based method to identify hub regions participating in between-network integration stratified by arousal, by measuring *k*-hubness, the number (*k*) of functionally overlapping networks in each brain region. We show evidence of a brain-wide decrease in between-network integration and inter-subject variability at low relative to high arousal, with differences emerging across regions of the frontoparietal, default mode, motor, limbic, and cerebellum networks. State-dependent changes in k-hubness relate to the actual patterns of network integration within these hubs, suggesting a brain state transition from high to low arousal characterized by global synchronization and reduced network overlaps. We demonstrate that arousal is not limited to specific brain areas known to be directly associated with arousal regulation, but instead has a brain-wide impact that involves high-level between-network communications.

## 1. Introduction

Fluctuations of brain and behavioral states during wakefulness are linked with our ability to observe and interact with a changing environment(Gonzalez-Castillo et al., 2021; McGinley et al., 2015; Zagha and McCormick, 2014). In the absence of external tasks, arousal, a behavioral state of being alert, awake, and attentive(Joshi and Gold, 2020; Liu and Falahpour, 2020), is a potential source of brain state variations or time-varying patterns of brain activity during resting state. Recent neuroimaging studies using functional magnetic resonance imaging (fMRI) show that there are neural correlates of arousal in the cerebral cortex(Breeden et al., 2017; DiNuzzo et al., 2019; Schneider et al., 2016; Yellin et al., 2015). In addition to key brain regions of the ascending arousal system, thalamo-cortical and cortico-cortical neural pathways are involved in modulation of arousal(Lee and Dan, 2012; McCormick et al., 2020; Paasonen et al., 2018), suggesting a role of arousal on functional interactions between brain regions. How arousal modulates functional brain organization during resting state remains poorly understood(Barttfeld et al., 2015; Shine et al., 2016; Yeo et al., 2015).

Functional connectivity is widely used to infer a relationship between brain regions by measuring the temporal correlation strength of the blood-oxygen-level-dependent (BOLD) signal. Integration of distinct brain regions can be described by connecting nodes (each representing a brain region) based on the strength of functional connectivity between them(Bullmore and Sporns, 2009). Hubs are defined as the nodes with a large number of connections to other nodes(Power et al., 2013). Among hubs, connector hubs play a key role in communications between networks, each being a set of inter-connected nodes. Connector hubs are thought to reconfigure their functional connectivity to adapt to changes in brain states such as tasks(Bertolero et al., 2015) or arousal(Boveroux et al., 2010). Brain-wide decreases in network integration were found in patients in the comatose state(Achard et al., 2012), under propofol-induced sedation(Qiu et al., 2017; Schrouff et al., 2011; Vatansever et al., 2020) and sleep(Boly et al., 2012). Still, the much more subtle question as to whether modulations in arousal during resting state are associated with brain-wide connector hub re-organization remains largely unexplored in healthy non-pharmacologically altered participants(Shine et al., 2016).

In addition, functional connectivity as measured by resting state fMRI involves complex information resulting from a variety of neurobiological, hemodynamic, and physiological components(Cole et al., 2014; Gonzalez-Castillo et al., 2019; Lurie et al., 2020). Components of time-varying functional connectivity at rest have been linked to consciousness(Barttfeld et al., 2015) and ongoing cognition(Gonzalez-Castillo et al., 2015). Other studies have observed time-varying resting state functional connectivity associated with sampling variability, motion artifacts, sleep states(Haimovici et al., 2017; Laumann et al., 2017), physiological noise(Chang et al., 2013), neurovascular coupling(Archila-Meléndez et al., 2020), or eye movements(Chang et al., 2016; Koba et al., 2021). How changes in arousal are linked to functional connectivity reconfiguration is a key question in understanding connectivity dynamics and could contribute to the heterogeneity of resting state functional connectivity patterns across subjects(Barttfeld et al., 2015; Laumann et al., 2017; Liu and Falahpour, 2020; Shine et al., 2016). However, level of arousal is not routinely monitored in most resting state fMRI studies, making it challenging to explore its impact on functional connectivity.

In this study, we identify and quantify changes in network integration stratified by arousal and examine its contribution to inter-subject functional connectivity variability. We hypothesize that resting state networks reorganize with arousal fluctuations. Specifically, we expect network integration to be lower at low relative to high arousal. To test this hypothesis, we collected resting state fMRI simultaneously with in-scanner pupillometry data from 27 healthy participants, in order to use pupillometry as an index of arousal(Larsen and Waters, 2018; Murphy et al., 2014; Schneider et al., 2016). We stratify high and low arousal states by ranked pupil area and estimate connector hubs in each state, using a recently introduced method; SParse dictionary learning based Analysis of Reliable K-hubness (SPARK)(Lee et al., 2018; Lee et al., 2016). SPARK identifies a set of individually consistent networks and defines connector hubs by measuring “*k*-hubness”, or the number (*k*) of functionally overlapping networks for each node(Lee et al., 2018; Lee et al., 2016). We show evidence of a brain-wide decrease in network integration and inter-subject variability of connector hubs in low versus high arousal resting states. By studying the hierarchical network organization of connector hubs, we observe that arousal is not localized to specific brain areas known to be directly associated with arousal regulation, but instead has a more extensive, brain-wide impact that involves high-level between-network communications.

## 2. Materials and Methods

### 2.1. Participants

This study was approved by the Institutional Review Board at Yale University. We recruited 37 healthy young adults (26.68 ± 4.18 years old; 20/17 females/males; 35/2 right/left-handed. Mean ± standard deviation) from the community of Yale University. Participants had to meet the following inclusion criteria: i) no claustrophobia or ferromagnetic metal in the body, ii) no clinical diagnosis of cognitive or mental disorders, iii) no visual impairments or difficulty in vision without glasses or contact lenses, and iv) no auditory impairments. Subjects were instructed to have a normal sleep before the day of scan and reported 7 ± 1 hours of sleep during the past 24 hours prior to the scan, with a neutral sleep quality scoring 3.4 ± 0.8 out of the five self-rating items: 1 (very bad), 2 (fairly bad), 3 (neutral), 4 (fairly good), 5(very good). Subjects reported a mild level of fatigue scoring 1 ± 0.7 out of the five self-rating items: 4 (worst possible fatigue) 3 (severe fatigue) 2 (moderate fatigue) 1 (mild fatigue) 0 (energetic, no fatigue). After data preprocessing, 10 subjects were excluded based on the following criteria: i) motion, estimated as the mean frame-to-frame displacement (FFD) > 0.15 mm in either of two resting state fMRI runs(Horien et al., 2019), ii) more than 35% of missing datapoints in the pupillometry data, or iii) missing data due to technical problems. Given these criteria, we included 27 subjects (26.52 ± 4.04 years old; 16/11 females/males; 25/2 right/left-handed) in our analyses. The mean FFD was 0.06 ± 0.02 mm for rest 1 and 0.06 ± 0.02 mm for rest 2 across the finally selected 27 subjects. Across the 27 subjects, the percent of discarded time-points was 6.02 ± 9.48 % for rest 1 and 4.33 ± 8.33 % for rest 2. See Table S1 for demographics.

### 2.2. Data acquisition

Imaging data were acquired using a Siemens 3.0T MAGNETOM Prisma MRI scanner at the Yale Magnetic Resonance Research Center. T1-weighted anatomical images were acquired using a magnetization prepared rapid gradient echo (MPRAGE) pulse sequence with the following parameters: repetition time (TR) = 2,400 ms, echo time (TE) = 1.22 ms, flip angle = 8°, slice thickness = 1 mm, in-plane resolution = 1 × 1 mm. Functional T2*-weighted BOLD images were acquired using a multiband gradient echo-planar imaging (EPI) pulse sequence (TR = 1000 ms, TE = 30 ms, flip angle = 55°, multiband acceleration factor = 5, slice thickness = 2 mm, 75 contiguous slices). Total duration of each functional run was 6:50 min (410 frames). Eye-tracking data were recorded using a MR-compatible infrared EyeLink 1000 Plus eye-tracking system (SR Research Ltd. Ottawa, ON, Canada) to measure time-varying changes in pupil area with a sampling rate of 1,000 Hz.

### 2.3. Pupillometry data preprocessing

Eye-tracking data were preprocessed using custom code in MATLAB R2018a. Eye blinks were automatically identified by EyeLink tracker’s online parser. Blink-induced artifacts were corrected using 4-point spline interpolation(Mathôt et al., 2013). Blinks that occurred shortly after each other (< 100 ms) were combined and treated as a single blink(Schneider et al., 2016). The signals were low-pass filtered using a first-order Butterworth filter at cut-off 0.5 Hz, after which the first 10,000 data points (i.e., 10 s) were removed to synch with the fMRI data. The time-course was down-sampled by averaging 1,000 consecutive frames for each 1 s bin, to match the fMRI sampling frequency (1 Hz). Pupil area was z-transformed using the mean and standard deviation over the rest 1 and 2, to control for variability in average pupil area across subjects. To account for the slow response time of the pupil to neuronal activity(Schneider et al., 2016), each time-course was convolved with the canonical hemodynamic response function (HRF) generated based on the mixture of two Gamma functions using SPM8(Friston et al., 1998). Finally, the normalized pupil area time-courses from rest 1 and 2 were temporally concatenated for each subject, to match the concatenated rest fMRI scans. We quantified eye-closure related missing pupillometry data by the proportion of missing (zero-valued) time-points with respect to the total number of time-points.

### 2.4. fMRI data preprocessing

T1-weighted anatomical images were skull-stripped using FSL optiBET(Lutkenhoff et al., 2014). All further analyses were performed using *BioImage Suite* unless otherwise specified(Joshi et al., 2011). Skull-stripped anatomical images were non-linearly registered to the standard Montreal Neurological Institute (MNI) space(Scheinost et al., 2017). Functional images were first motion-corrected and realigned using twenty-four motion parameters(Satterthwaite et al., 2013), including six rigid-body parameters, their temporal derivatives, and their quadratic terms, using SPM8. Subject within scan head motion was quantified by computing the mean FFD across each functional run. The first 10 s volumes were discarded to exclude frames when eye-tracking system was initialized and stabilized. Functional images were linearly registered to skull-stripped anatomical images using the rigid transformation of the mean functional image from the first run (rest 1). 3D spatial smoothing was performed using an isotropic Gaussian kernel with a 4 mm full-width-half-maximum(Scheinost et al., 2014). Nuisance covariates, including 1) 24 motion parameters, 2) slow temporal drifts as modeled by the linear, quadratic, and cubic Legendre polynomials, 3) the mean signals in the cerebrospinal fluid and white matter, and 4) the whole-brain global signal, were regressed. Data were low-pass filtered using a zero-mean unit-variance Gaussian filter with a cut-off frequency of 0.12 Hz. For each run, using a 268-parcel functional atlas that covered the whole brain excluding ventricles and white matter(Finn et al., 2015; Shen et al., 2013), we generated a time-by-node data matrix for each individual by averaging the fMRI signals across voxels within each node. Finally, two data matrices from rest 1 and 2 were temporally concatenated for each subject.

### 2.5. Hub analysis

An overview of our analysis pipeline is shown in Fig. 1. We use the pupillometry data, ranked by pupil area, to estimate arousal at each time-point. We stratify high and low arousal states by selecting time-points from the top 20% (large) and bottom 20% (small) pupil areas, resulting in 160 frames of fMRI data for each state. Using data from either the high or low arousal state, we estimated resting state networks and nodal *k*-hubness using SPARK (Lee et al., 2019; Lee et al., 2016). SPARK represents the BOLD signals from each node as a sparse, weighted linear combination of atoms, where each atom is a temporal feature of each network(Lee et al., 2011). When there are subject-specific *N* networks in the whole brain, a node involves a node-specific combination of *k* (less than *N*, therefore, sparse) networks, identifying temporally and spatially overlapping networks. *k*-hubness is a measure of between-network integration and can serve to identify connector hubs, which are obtained by counting the number of functional networks that overlap at each node.

**Figure 1.**
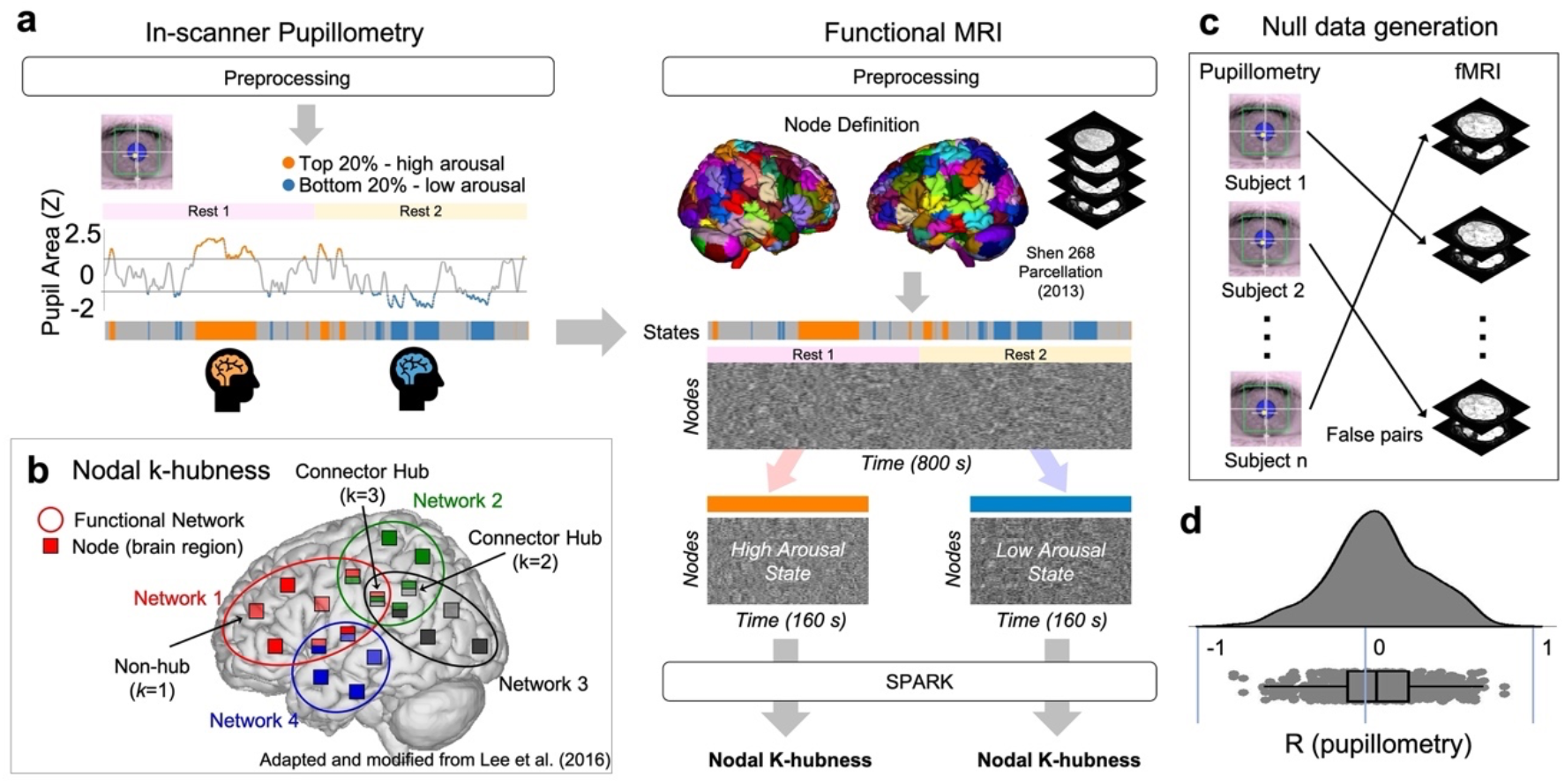
Analysis pipeline overview. (**a**) Pupillometry-based fMRI state stratification for arousal level-dependent connector hub analysis. For each subject, pupillometry data were used to stratify the simultaneously acquired fMRI data into two states (high and low arousal). Specifically, time-points where pupil area was within the top or bottom 20% rank were assigned to a high-(orange) or low-arousal state (blue), respectively. A sparsity-based analysis of reliable *k*-hubness (SPARK) was used to identify connector hubs from state-stratified fMRI data, by measuring *k*-hubness for each node at the individual level. (**b**) *k*-hubness is defined as the number of overlapping networks in each node. (**c**) Null data generation by randomizing the assignment of pupillometry to fMRI across the 27 subjects. (**d**) The distribution of Pearson’s correlation coefficients measured between individual pupillometry time-courses.

In this work, SPARK was applied for an individual fMRI data (a time by node matrix) as follows (Fig. S1 for a summary diagram). First, a circular block bootstrap with a block length *h* was performed to generate 300 surrogate datasets with equal dimensions(Bellec et al., 2010). Using a proper choice of *h* (e.g. equal to or larger than the square root of the number of time-points) to preserve temporal structures in the BOLD signal, we assume that resampled data are from the same probabilistic distribution(Bellec et al., 2010). SPARK aims to detect highly consistent hubs across a large number of bootstraps. For each resampled dataset, a sparse dictionary learning algorithm (Aharon et al., 2006; Lee et al., 2018; Lee et al., 2016) was applied to learn a dictionary involving *N* time-course atoms (temporal features) and a corresponding sparse coefficient matrix (spatial maps). The algorithm involves an automatic parameter estimation strategy using the minimum description length criteria(Lee et al., 2018). The total number of networks (*N*) was estimated independently for each resampled dataset by varying *N* from 1 to the number of principal components that explained 99% of the variance in each resampled dataset. The level of sparsity (*k*) was determined by varying *k* from 1 to *N/2* for each *N*. The reproducibility of parameters (*N* and *k*) of the sparse model were assessed across bootstraps.

After the 300 parallel processes, we collected 300 sparse coefficient matrices and applied K-means spatial clustering. The number of clusters was the median of estimated *N* across 300 resampled datasets. The spatial maps were averaged within each cluster, and thresholded at 95% confidence interval by approximating the Gaussian distribution of background noise in the average matrix. This provided the final sparse matrix, in which each row represented a spatial map of individually reliable resting state networks. Counting the non-zeros for each column of this matrix provided an estimation of *k*-hubness for each node. This clustering procedure was repeated 100 times to take into account random initializations in K-mean clustering. Finally, nodal *k*-hubness was determined by the mean of *k*-hubness values estimated over 100 clustering results. The density of *k*-hubness was calculated as the % proportion of nodes with non-zero *k*-hubness to the total number of nodes estimated from data obtained in each arousal.

For comparison purposes, we generate null data by randomizing the assignment of pupillometry to fMRI across the 27 subjects (Fig. 1c). This results in 702 false fMRI-pupillometry pairs, from which we stratify random high/low arousal assignments. Pupillometry time-courses are unique to the individual (Fig. 1d). The distribution of pupil time-course correlations is not skewed (skewness= −.04) and not normal (Lilliefors’ test, *p*<.003). See Fig. S2 for the individual pupillometry time-courses. We compare our results to those from null data, in order to test the null hypothesis that there is no association between resting state functional connectivity and spontaneous arousal fluctuations defined using pupillometry.

### 2.6. Hub disruption index

To assess the brain-wide connector hub reorganization with arousal, we defined the hub disruption index (HDI_k_) using *k*-hubness from resting state fMRI at high and low arousal states(Lee et al., 2018). The HDI was first proposed for studying hubs defined using degree centrality in graph theory(Achard et al., 2012) and introduced for *k*-hubness to study the reorganization of connector hubs in patients with epilepsy(Lee et al., 2018). The HDI_k_ is a summary measure to quantify overall hub reorganization across the whole brain between control (e.g., high arousal) and experimental (e.g., low arousal) brain states(Achard et al., 2012). We measured the HDI_k_ at both the group and single subject levels. At the group level, HDI_<k>_ is the slope of the linear regression model fit to group average *k*-hubness across subjects at high arousal (*x*-axis, *<k>*_High_) and the difference in group average *k*-hubness between low and high arousal (*y*-axis, *<k>*_Low_ - *<k>*_High_) (Fig. 2a). A negative slope means that some of the hubs identified at high arousal lose their hub status at low arousal (i.e., nodes exhibiting a high *<k>*_High_ relative to *<k>*_Low_ have a negative value of *<k>*_Low_ - *<k>*_High_). HDI_<k>_ becomes zero when there is no difference in *k*-hubness between the two states. The same approach is used to define HDI_k_ at the individual level, by using the individual subject’s nodal *k*-hubness (*k*) in a single subject (*x*-axis, *k*_High_; *y*-axis, *k*_Low_ - *k*_High_).

**Figure 2.**
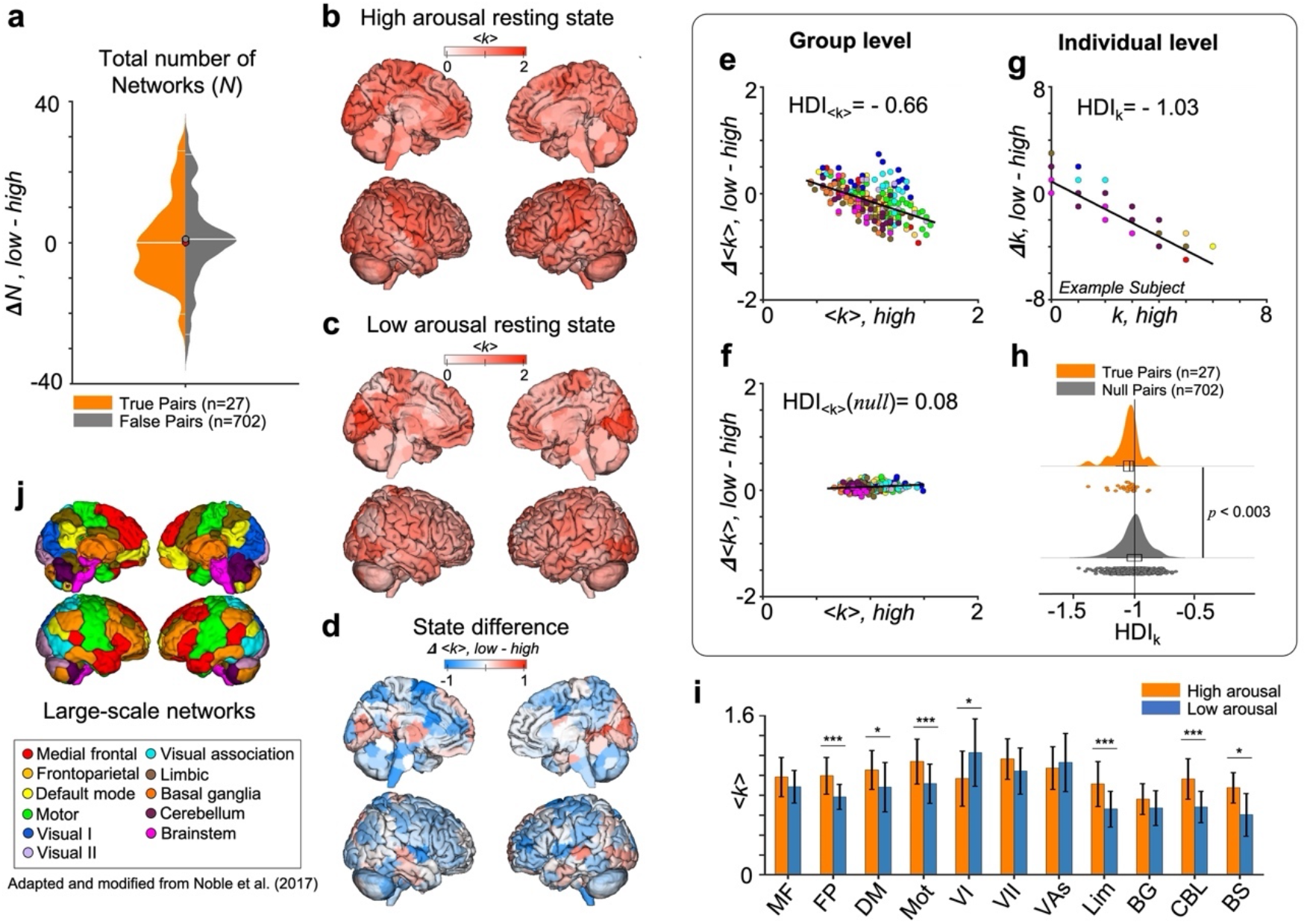
Distributed connector hubs are re-organized with arousal modulations during resting state. (**a**) The total number of resting state networks (*N*) detected by SPARK from individuals were preserved between high and low arousal states. (**b-c**) The group average *k*-hubness maps at high (b) and low (c) arousal. (**d**) The map of difference in the group average *k*-hubness between the low and high arousal states. (**e**) The estimation of group-level HDI_<k>_ between high and low arousal. A linear regression model is used to find a linear fit of nodal group-average *k*-hubness (<k>) estimated from the two states. HDI_<k>_ is defined as a slope of the linear fit. (**f**). The estimation of group-level HDI_<k>_ between two randomized states, by averaging *k*-hubness across 702 false brain-pupil pairs in each node. (**g**) An example of individual-level HDI_k_ from a single subject exhibiting the median of HDI_k_ within group. Note that nodal *k*-hubness is an integer, therefore nodes with a same value are superimposed in this scatter plot. (**h**) The distribution of individual-level HDI_k_ (top) and those from null data (bottom). *p*-value estimated using the left-tailed Wilcoxon rank sum test is shown. (**i**) The bar plot of *k*-hubness distributions within the eleven pre-defined large-scale networks in each state. Mean ± standard deviation. (**j**) Data points in figures e-f and j are color-coded using eleven *a priori* functional networks. Ten networks were defined as described in Noble et al.(Noble et al., 2017), and the nodes belonging to the brainstem were assigned to an 11th network.

### 2.7. Hub connectivity probability

To investigate whether and how state-dependent changes in connector hubs relate to the actual patterns of network integration within these hubs, we computed the conditional probability (***p***_*i*_) of each node *i* to be a member of functional networks overlapping in a hub *j.* To do this, for each arousal state, we first collected all resting state network maps estimated from all individual subjects. Using this collection, ***p***_*i*_ was computed by the proportion of the number of functional networks involving both a node *i* and the hub *j* over all subjects to the total number of networks that involved the node *j* over all subjects, such that ***p***_*i*_ =1 if *i* = *j*.

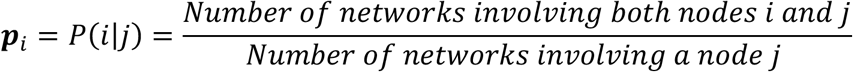

This provided a probability map of functional connectivity associated with a hub *j*. A high probability ***p***_*i*_ indicates that a node *i* is more likely to be a part of functional connectivity associated with a specific hub *j* across subjects, or the extent to which a connector hub contributes to inter-subject consistency of functional connectivity integration across the brain. Next, we calculated for each node *i* the total functional connectivity across the whole brain as the total probability ***P***_*i*_:

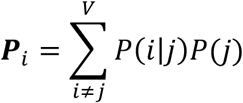

where *P*(*j*) = 1/(*V* − 1) and *V* is the total number of nodes in the brain. The total probability ***P***_*i*_ indicates the amount of nodal functional connectivity associated with distributed hubs over the whole brain. Note that the total number of networks involving a hub is state dependent; therefore, it is possible to normalize within state. Then, a transition vector was identified for each node within the scatter plot of the group average *k*-hubness (*<k>*, x-axis) and the total probability (***P***_*i*_, y-axis), as a vector that links a node at high arousal state (<k>_high_, ***P***_*i*(high)_) to the same node at low arousal state (<k>_low_, ***P***_*i*(low)_). To visualize the magnitude and direction of transition vectors for all nodes, the transition vectors were re-centered to have a link from (0,0) to (<k>_low_- <k>_high_, ***P***_*i*(low)_-***P***_*i*(high)_).

## 3. Results

We present the results from our connector hub analyses across arousal levels as follows. First, we assessed whether the global scale of functional connectivity is preserved across high and low arousal states (Section 3.1). We then estimated hub disruptions in the whole brain at both the group and single subject levels (Section 3.2). Next, we investigated the impact of arousal on connector hubs in large-scale networks (Section 3.3) and inter-subject variability of the connector hub organizations (Section 3.4). Then, we studied whether and how such connector hub disruptions relate to the actual patterns of functional network integration within these hubs (Section 3.5). Lastly, the reliability and robustness of our hub estimations are addressed in Section 3.6.

### 3.1. Preserved global network scale between high and low arousal states

We first assessed whether the total number of functional networks in the whole brain is preserved across high and low arousal states. To avoid any potential confounds introduced by the state stratification strategy in our parameter estimations (e.g., the number and duration of continuous state segments), we did not directly compare the distributions of *N* between high and low arousal states. Instead, we compared the between-state differences in *N* to the difference observed from null data (Wilcoxon rank sum test, *p*>.5). In-line with previous work(Achard et al., 2012; Vatansever et al., 2020), we found that the total number of networks (*N*) detected by SPARK from individuals was preserved between states (Fig. 2a). Estimated *N* was 30 ± 16.3 (median ± interquartile range) at high arousal and 25 ± 8 at low arousal. The goal is to investigate whether the patterns of connector hubs actually change with arousal modulations, while the global network scale is preserved during resting state.

### 3.2. Brain-wide disruptions of connector hubs from high to low arousal states

We observed differences between the group average *k*-hubness maps estimated from resting state fMRI at high and low arousal states. At high arousal, connector hubs are widely distributed across the unimodal and transmodal cortices, subcortical structures, cerebellum and brainstem (Fig.2b). At low arousal, we observe an overall decrease in *k*-hubness across the brain, relative to the high arousal state, with the exception of some nodes in the visual networks (Fig. 2c-d, Fig. 2i, two-sided Wilcoxon rank sum test, Bonferroni corrected *p*<.05).

To quantify the overall degree of hub disruptions in the whole brain, we defined the hub disruption index (HDI_k_) using *k*-hubness (Lee et al., 2018). We found the group-level HDI_<k>_ to be −0.66, indicating a brain-wide disruption of connector hubs with arousal at resting state (Fig. 2e). From the null data, the estimated HDI_<k>_ was 0.08, indicating no hub disruption between randomly stratified states (Fig. 2f). We next assessed if our group-level finding was replicated in individual subjects. To do this, using the same approach, we define HDI_k_ using the individual subject’s nodal *k*-hubness (*k*) to quantify the overall connector hub reorganization at the individual level (*x*-axis, *k*_High_; *y*-axis, *k*_Low_ - *k*_High_) (Fig. 2g). In Fig. 2h, the distribution of individual level HDI_k_ estimated from 27 subjects (HDI_k_= −1.03 ± 0.08) is shown compared to that from 702 randomized pupillometry samples (HDI_k_ = −0.99 ± 0.12). Consistent with the group results, we observed a negative relationship indicating that connector hubs reorganize from high to low arousal at the individual subject level, when compared to the results from null data (left-tailed Wilcoxon rank sum test, *p*<.003) (see Fig. S3 for the individual level results from all subjects). The group- and individual-level HDI analyses for *k*-hubness demonstrate arousal-level-dependent changes in between-network integration in resting state functional connectivity.

Motion can be a difficult confound in fMRI analyses(Power et al., 2015; Satterthwaite et al., 2012). We assessed if individual-level hub disruption was correlated with subject motion in fMRI data and if so, would it account for these findings. We show that inter-subject variability of the estimated HDIk is not correlated with head motion (Wilcoxon rank sum test, p>.8) (Fig. S4). We also tested if individual-level hub disruption was correlated with the proportion of missing data-points in the pupillometry data. The proportion of missing data-points may affect identification of the arousal state. Potential causes of missing data included technical errors such as a connection issue with the eye-tracking system and inability to quantify pupil size due to blinks and saccades. Across the 27 subjects, we found 6.02 ± 9.48 % discarded time-points for rest 1 and 4.33 ± 8.33 % for rest 2 (Table S1). Inter-subject variability of the estimated HDIk was not correlated with missing data (Wilcoxon rank sum test, p>.2) (Fig. S5).

### 3.3. Brain-wide decreases in network integration at low relative to high arousal

We next investigated the impact of arousal on large-scale networks. In Fig. 3a, we show the distribution of arousal-level-dependent changes in group-average *k*-hubness (Δ*<k>, low - high*) within each large-scale network(Noble et al., 2017). For each network, we compare the Δ*<k>* distribution from the nodes belonging to this network estimated across our 27 subjects (color-coded), to the null Δ*<k>* distribution from the same nodes estimated from randomized data over 5,000 permutations (in gray color). To do this, the null Δ*<k>* distribution was obtained by averaging nodal *k*-hubness across the same 27 subjects with a randomized set of brain-pupillometry pairs. As a result, we found decreases in group-average *k*-hubness with decreased arousal in the frontoparietal, motor, limbic and cerebellum networks (two-tailed Wilcoxon rank sum test, Bonferroni corrected *p*<.001) and the default mode network (*p*<.01).

**Figure 3.**
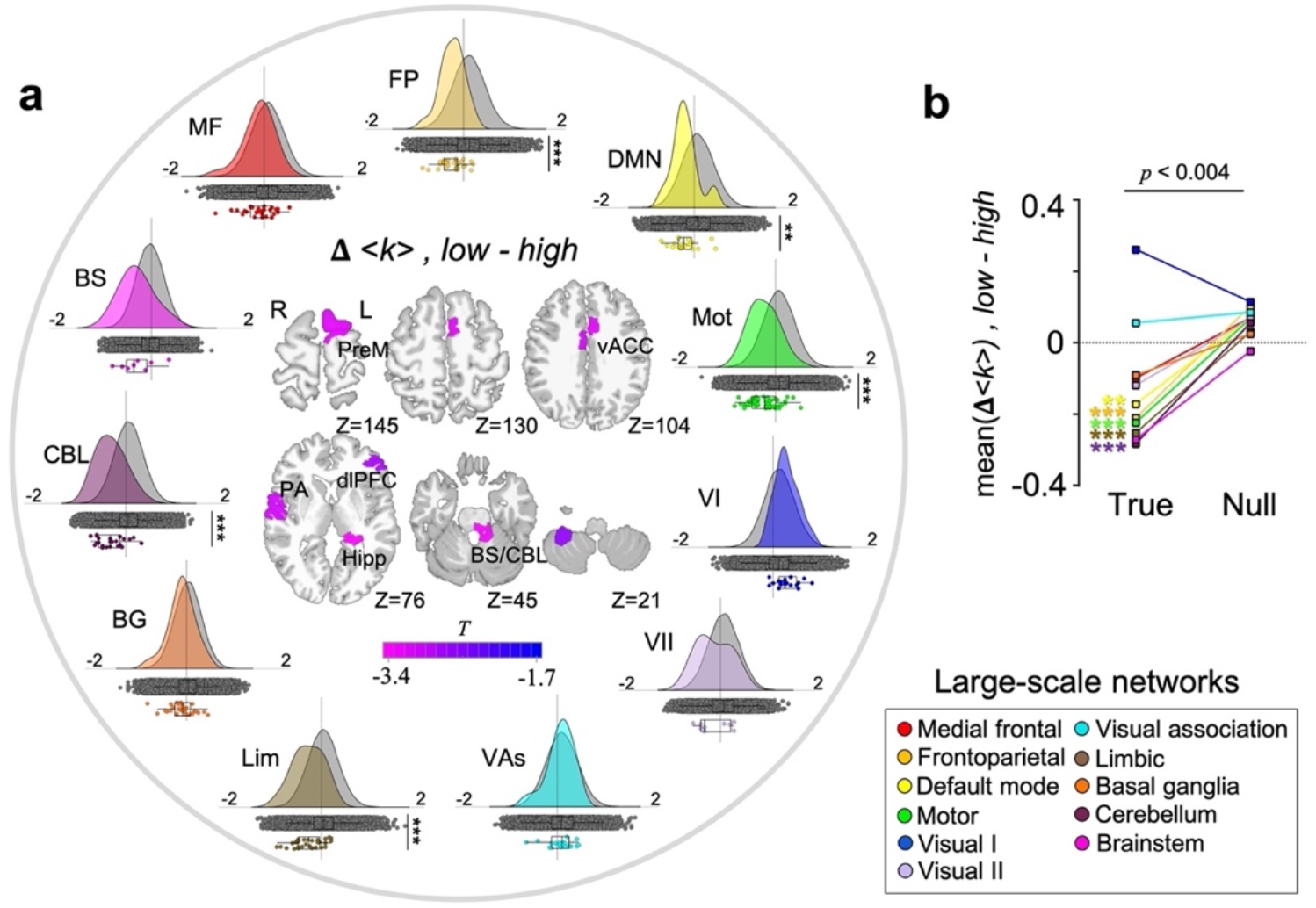
Decreased network integration at low relative to high arousal. (**a**) Around the circle, we show the distributions of between-state changes in group-average *k*-hubness (Δ*<k>, low-high*) within each of the pre-defined large-scale networks (color-coded). The null distribution of Δ*<k>* was generated from the same nodes in each network over 5,000 permutations (shown in grey). Asterisks indicate Bonferroni corrected *p*-values from the two-tailed Wilcoxon rank sum tests. At the center of the circle, we show a node-wise two-sample test result (one-tailed bootstrap test, FDR corrected p<.05) with 5,000 bootstraps(Efron and Tibshirani, 1994). (**b**) A summary of network-level Δ*<k>* distributions that are shown in (a), using the mean of Δ*<k>* within each network. *p*<.004 using the two-tailed Wilcoxon rank sum test. PreM: Premotor cortex. vACC: ventral anterior cingulate cortex. PA: primary auditory cortex. dlPFC: dorsolateral prefrontal cortex. Hipp: hippocampus. BS/CBL: brainstem/cerebellum.

In addition, node-wise statistical tests on individual connector hubs confirmed our observation of a brain-wide decrease in between-network integration, and highlighted nodes that exhibited consistent changes across subjects (Fig. 3a, at the center of circle plot). We used the left-tailed bootstrap-based two-sample tests proposed in Efron et al.(Efron and Tibshirani, 1994), because individual-level *k*-hubness is a discrete integer within a small range (e.g. [0, 5]) and the symmetry of distributions are not assumed. We found decreases in *k*-hubness at low arousal in the premotor/supplementary motor, ventral anterior cingulate, primary auditory and dorsolateral prefrontal cortices, hippocampus, cerebellum, and in the node that spans from the cerebellum to the locus coeruleus in the brainstem (Z= −24 in the MNI coordinates)(Keren et al., 2009) (FDR corrected *p*<.05, see Table S2). In Fig. 3b, we summarize our findings using the mean of Δ*<k>* within each network, highlighting arousal-dependent decreases in group-average *k*-hubness in the five large-scale networks.

### 3.4. Inter-subject variance in network integration decreases from high to low arousal

Next, we quantified the change in inter-subject variance of nodal *k*-hubness between arousal states. Figure 4a illustrates the map of between-state differences in the standard deviation of *k*-hubness (*σ*_*k*_). At low relative to high arousal, we found a brain-wide decrease in inter-subject variance of *k*-hubness across the brain in regions belonging to the medial frontal, frontoparietal, default mode, motor, limbic, cerebellum and brainstem networks (Fig. 4b, two-sided Wilcoxon rank sum test, Bonferroni corrected *p*<.05). To test if such decreases were above chance, we computed the between-state difference in standard deviation of *k*-hubness (Δ*σ*_*k*_, low – high) across 27 subjects. The null Δ*σ*_*k*_ distribution was obtained by averaging nodal *k*-hubness across the same 27 subjects with a randomized set of brain-pupillometry pairs. Figure 4c shows that the distribution of Δ*σ*_*k*_ from the 268 nodes in the whole brain is lower than the null Δ*σ*_*k*_ distribution (two-sided Wilcoxon rank sum test, *p*<4e-23). In each pre-defined large-scale network, we compared the Δ*σ*_*k*_ distribution from the nodes belonging to each network estimated from 27 subjects to the null Δ*σ*_*k*_ distribution (in gray color). In Fig. 4d, we summarize our findings using the mean of Δ*σ*_*k*_ within each network. We found decreases in inter-subject variance of *k*-hubness with arousal, again, in the frontoparietal (two-sided Wilcoxon rank sum test, Bonferroni corrected *p* <.05), default mode (*p*<.01), motor, limbic and cerebellum networks (*p*<.001). We did not find state-differences in inter-subject variability at the node level (Levene’s test for equality of variance, 5,000 permutations, FDR corrected *p*<.05).

**Figure 4.**
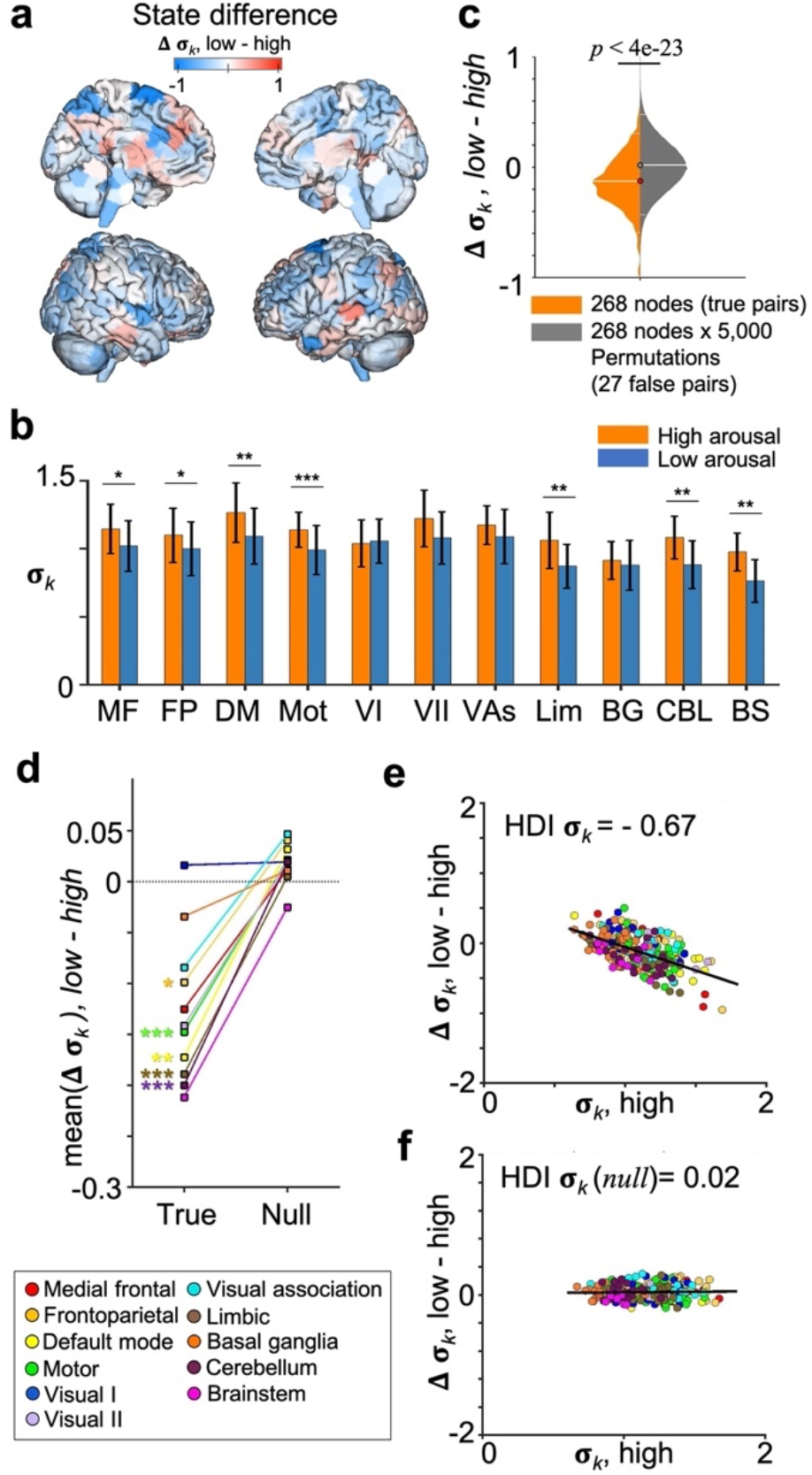
Inter-subject variability in functional network integration decreases from high to low arousal. (**A**) The map of between-state difference in the standard deviation of *k*-hubness (*σ*_*k*_) across subjects; between low and high arousal resting states. (**B**) The bar plot of *σ*_*k*_ distributions within the eleven pre-defined large-scale networks in high and low arousal states. Mean ± standard deviation. (**C**) The distribution of between-state differences in inter-subject variance of *k*-hubness (Δ*σ*_*k*_, low – high), estimated across 27 subjects. The null distribution of Δ*σ*_*k*_ was generated over 5,000 permutations. (**D**) A summary of network-level Δ*σ*_*k*_ distributions using the mean of Δ*σ*_*k*_ within each network (left) compared to the null distribution of Δ*σ*_*k*_ (right). (**E**) Brain-wide changes in *σ*_*k*_ between high and low arousal. The hub disruption index (HDI) for nodal *σ*_*k*_ reveals a brain-wide decrease in inter-subject variability. (**F**) Whereas there is no difference observed for null data. Asterisk indicates statistical significance from Wilcoxon rank sum tests with Bonferroni corrected *p*-values, *: *p*<.05, **: *p*<.01, ***: *p*<.001.

To quantify brain-wide changes in inter-subject variability between low and high arousal, we defined a HDI for *σ*_*k*_, using the method to estimate the HDI for *k*-hubness. We found a negative slope (−0.67), indicating a brain-wide decrease in inter-subject variability at low relative to high arousal (Fig. 4e). For null data, the slope was 0.02, which is close to zero, indicating no difference in state-dependent HDI for k-hubness (Fig. 4f). Note that the range of *σ*_*k*_ for real and null data in Fig. 4e/4f (e.g., less than 2), indicating that there is no sample size bias. Taken together, our results demonstrate an overall decrease in inter-subject variability of *k*-hubness comparing the low relative to high arousal state.

### 3.5. Resting state networks at low arousal have reduced network overlaps and increased global connectivity

It is important to investigate how such connector hub disruptions relate to the actual patterns of functional network integration within these hubs. For each arousal state, we generated a probability map of functional connectivity involving each hub, by computing the conditional probability (***p***_*i*_) of each node *i* to be a member of any functional network overlapping in a hub *j* (Fig. 5a). This conditional probability was computed from the pooled collection of all networks estimated from all individual subjects. The total number of hub-related (pooled) overlapping networks was 32 ± 11 (median ± interquartile range across the 268 nodes) at high arousal and 24 ± 11 at low arousal (Fig. 5b; two-sided Wilcoxon rank sum test, *p*<2e-26). This indicates that there are fewer network overlaps resulting in lower between-network integration at low arousal. In addition, the spatial distribution of ***p***_*i*_ varied across hubs. For example (Fig. 5c), for a connector hub in the right ventral anterior cingulate cortex, the spatial distribution of functional connectivity integrated with this hub was broader in the low arousal, suggesting lower inter-subject variance at low relative to high arousal. On the other hand, the probability map for a connector hub in the left dorsolateral prefrontal cortex shows a similar spatial distribution of hub-associated functional connectivity at both high and low arousal, including regions of the frontoparietal and default mode networks, suggesting stable inter-subject variability across arousal levels. Next, we quantified the total amount of functional connectivity of each node *i* over the whole-brain as the total probability ***P***_*i*_ (Fig. 5d). The median of ***P***_*i*_ distribution was higher at low arousal relative to high arousal (two-sided Wilcoxon rank sum test, *p*<2e-26), suggesting an increase in global connectivity. A scatter plot of the total probability (***P***_*i*_) and the group average *k*-hubness (*<k>*) estimated for 268 nodes shows a clear pattern of connector hub disruption: a decrease in *k*-hubness and an increase in ***P***_*i*_ from high to low arousal (Fig. 5e). In addition, as expected, we found that nodes exhibiting large group-average changes in <k> also exhibit large changes in inter-subject variability (**g**) and total connectivity probability (**h**) (rs: Spearman’s rank correlation, *p*=0).

**Figure 5.**
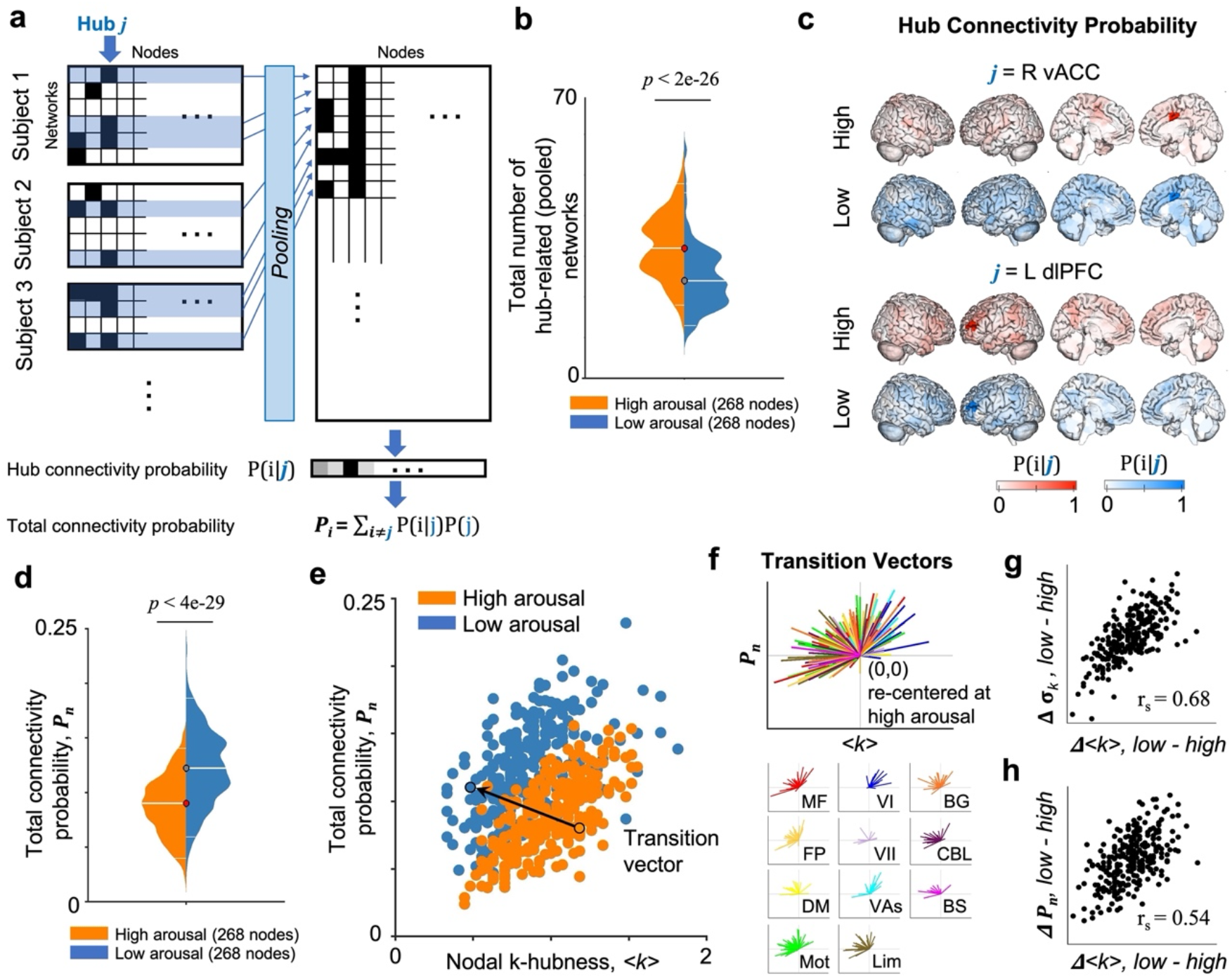
Resting state networks at low arousal have reduced network overlaps while exhibiting brain-wide connectivity. (**a**) A summary diagram to calculate hub connectivity probability (***p***_*i*_) and total connectivity probability (***P***_*i*_). For an arousal state, resting state networks involving a hub *j* are collected from all subjects. ***p***_*i*_: the conditional probability of each node *i* to be a member of functional networks overlapping in a hub *j.* (**b**) The total number of hub-related networks for each node is lower at low relative to high arousal. (**c**) Probability maps of functional connectivity integrated in a specific hub (two exemplary nodes in the right vACC and the left dlPFC) across subjects, at high (red) and low (blue) arousal. (**d**) Total connectivity probability (***P***_*i*_) is higher at low relative to high arousal across the whole brain, indicating an increased global synchronization. (**e**) Scatter plot of hub measures from 268 nodes at high (orange) and low (blue) arousal data. x-axis denotes the group average *k*-hubness (*<k*>). y-axis denotes the total probability ***P***_*i*_ of hub functional connectivity (FC) calculated for each node *i*. Left (L)/Right (R) in bold. An exemplary transition vector that links a node at high arousal state (<k>_high_, ***P***_*i*(high)_) to the same node at low arousal state (<k>_low_, ***P***_*i*(low)_) is shown. (**f**) Re-centered transition vectors for all nodes, from (0,0) to (<k>_low_-<k>_high_, ***P***_*i*(low)_-***P***_*i*(high)_), show a trend pointing toward the quadrant II, indicating a decrease in *k*-hubness and an increase in ***P***_*i*_ from high to low arousal. Transition vectors for nodes in each large-scale network (color-coded as in Fig. 2-4) are shown below. Nodes exhibiting large group-average changes in <k> also exhibit large changes in inter-subject variability (**g**) and total connectivity probability (**h**) (rs: Spearman’s rank correlation, *p*=0).

### 3.6. Reliability and robustness

The reliability of hub estimations at the single subject level is assessed using SPARK(Lee et al., 2016), to extract the most reproducible patterns of overlapping functional networks. Within this procedure, we select highly reproducible components at 95% confidence interval (CI) by approximating the Gaussian distribution of background noise in network maps estimated from an average across 300 bootstraps. In this study, the density of *k*-hubness estimated from data obtained in the high arousal state, for instance, decreases with threshold: 0.84 ± 0.38 (median ± interquartile range) at 90% CI, 0.58 ± 0.41 at 95% CI, and 0.35 ± 0.29 at 99% CI (Fig. S6). This means only 35% of nodes are reliably involved in at least one network, when using the most conservative threshold. To validate whether our findings are robust to the choice of CI, we repeated our analyses using 90% and 99% CI. As expected, between-state changes in global network scale (Δ*N*, low-high) were preserved across arousal states. Furthermore, we observed decreases in the group-average *k*-hubness (Δ<k>) and their inter-subject variability (Δ*σ*_*k*_) at low relative to high arousal across all CI thresholds, with such changes being most robust in the motor and cerebellum networks (Fig. S6).

## 4. Discussion

Using simultaneous fMRI and pupillometry, we demonstrated brain-wide changes in network integration associated with fluctuations in arousal during the resting-state. We found decreases in between-network integration from high to low arousal by analyzing *k*-hubness at both the group- and individual-subject level. K-hubness differences emerged in regions including the frontoparietal, default mode, motor, limbic, and cerebellum networks. These findings establish a relationship between modulations in arousal during resting wakefulness and the dynamics of functional brain organization, including changes in connector hubs or between-network integration. The inter-subject variability of connector hubs decreases at low relative to high arousal, whereas the impact of arousal modulation on connector hub-related functional network integration differed between brain regions. State-dependent changes in connector hubs relate to the actual patterns of network integration within these hubs. While the global network scale, the total number of networks in the brain, was preserved between the high and low arousal states, the number of hub-related networks decreased, and the nodal total connectivity probability increased at low relative to high arousal state. These findings together suggest a brain state transition from high to low arousal characterized by global synchronization or reduced functional network specializations. Control analyses indicated that motion and eye-closure related effects are not driving results. Our results demonstrate that *k*-hubness is sensitive to arousal levels within resting state and that arousal is not localized to specific brain areas known to be directly associated with arousal regulation, but instead it’s associated with brain-wide changes involving high-level between-network communications.

These findings demonstrate the utility of simultaneous pupillometry as a proxy for measuring variations in arousal during resting-state fMRI. In the absence of task-related cognitive demands, pupil changes are mainly driven by non-specific factors such as arousal(Joshi and Gold, 2020; Liu and Falahpour, 2020). We observed brain-wide connector hub disruptions between low and high arousal, by measuring the hub disruption index of group-average *k*-hubness (HDI_<k>_) (Fig. 2). This finding indicates the flexibility of functional networks over time even during rest(Barttfeld et al., 2015; Shine et al., 2016; Yeo et al., 2015). The negative HDI_<k>_ at the group level was replicated using HDI_k_ values estimated from individual subjects. These results are in agreement with previous work using HDI for degree centrality in graph theory, reporting hub disruptions in patients with coma(Achard et al., 2012), and in healthy subjects with propofol-induced sedation(Vatansever et al., 2020). In this work, we were able to address a more subtle question as to whether modulations in arousal during the resting state are associated with changes in the functional organization of the brain. On the other hand, we found that the HDI_*k*_ estimated from the randomized state datasets was higher at the individual level than that estimated at the group level, potentially reflecting the presence of other factors (e.g., ongoing thought, emotion) that could account for a portion of the inter-subject variation. Future work should aim to identify such factors to understand the relationship between these factors and hub configuration in the resting state.

We found decreases in *k*-hubness at low relative to high arousal in regions of the frontoparietal, default mode, motor, limbic and cerebellar networks (Fig. 3). These regions have been implicated in previous work that assessed co-fluctuations of resting state BOLD activity and simultaneous pupillometry(Breeden et al., 2017; DiNuzzo et al., 2019; Schneider et al., 2016; Yellin et al., 2015). Schneider et al. found a positive coupling of pupil dilation with BOLD activity in the salience and default mode networks, frontal and parietal areas, and a negative relationship between spontaneous pupil constrictions and BOLD activity in the visual and sensorimotor areas(Schneider et al., 2016). Modulations of the default mode network have been observed during sleep deprivation(De Havas et al., 2012; Gujar et al., 2010; Yeo et al., 2015) and light sleep(Boly et al., 2012; Larson-Prior et al., 2011; Spoormaker et al., 2010; Sämann et al., 2011). We found a between-state change in *k*-hubness in the node that spans from the cerebellum to the locus coeruleus in the brainstem (Z= −24 in the MNI coordinates)(Keren et al., 2009), a core region of the ascending arousal system(Lee and Dan, 2012), in agreement with Murphy et al. who found that pupil diameter covaries with BOLD activity in the locus coeruleus(Murphy et al., 2014). Here, we extend these previous studies by demonstrating that modulations of arousal are not limited to specific brain areas directly associated with the brain’s ascending arousal system, but instead involve brain-wide communication networks.

That we found arousal-level-dependent decreases in between-network integration in regions of the frontoparietal cortex, suggests a role of arousal modulations in baseline activity related to cognition. Decreases in functional connectivity in the frontoparietal network were found during propofol-induced loss of consciousness and sleep(Boly et al., 2012; Boveroux et al., 2010; Schrouff et al., 2011; Schröter et al., 2012). Shine et al.(Shine et al., 2016) proposed that resting state functional connectivity alternates between integrated and segregated network topologies, and demonstrated a positive relationship between pupil diameter and network integration within these regions. Those findings in large part harmonize with our work, but there are several key differences. Notably, they identified integrated or segregated “topological” states from data, while we identified high or low “behavioral” arousal states. Our approach did not take into account intermediate levels of arousal and potential transient variations in hubs, but instead focused on detecting the most reproducible and individually consistent hub features characterizing each arousal state. Together, these findings lend support to the theory that state-dependent changes in brain functional connectivity may be driven by ongoing alterations in ascending neuromodulatory input and global fluctuations in neural gain(Eldar et al., 2013; Shine et al., 2016).

We found decreases in inter-subject variability of *k*-hubness at low relative to high arousal (Fig. 4). The global network scale, the total number of networks estimated in the whole brain, was preserved between the high and low arousal states (Fig. 2a). The number of networks involving hubs was reduced in the low relative to high arousal state (Fig. 5b). The total functional connectivity increased over the whole brain at low arousal, despite the reduced number of hub-related networks, suggesting a brain state transition from high to low arousal characterized by global synchronization or reduced functional network specializations. In addition, the impact of arousal modulation on connector hubs differed between brain regions (Fig. 5). This suggests that accounting for arousal-dependent changes may help understand individual variability in functional connectivity and its association with behavior. Functional connectivity has been shown to be valuable in identifying individuals using patterns of brain functional connectivity (i.e., fingerprinting)(Finn et al., 2015), and in predictive models relating functional organization to behavior both under rest-(Finn et al., 2015; Shen et al., 2017) and task-conditions(Finn et al., 2017; Greene et al., 2018; Rosenberg et al., 2015; Rosenberg et al., 2016). Task conditions offer a controlled manipulation of brain state, in contrast to the unconstrained nature of resting state; therefore, it is likely that individual differences in task-relevant circuitry can be amplified to help predict related traits(Greene et al., 2018; Lowe et al., 2000). Functional connectivity estimated from higher arousal resting state may play a different role in predicting traits, particularly for some phenotypes associated with high-level functions. It has already been demonstrated that state manipulations can influence trait predictions(Finn et al., 2017) for example. Given this evidence of the cognitive relevance of resting state functional connectivity(Barttfeld et al., 2015; Gonzalez-Castillo et al., 2019) in developing predictive models of behavior, future work should incorporate the role of arousal.

It should be noted that data were processed identically in both the high and low arousal states and in the null data set. Therefore, our observations cannot be attributed to some methodological artifacts, such as dwell time difference. The fMRI in the high and low arousal states, was balanced in terms of the amount of data included. We did not take into account the potential impact of other potential confounds, such as caffeine and alcohol consumption, anxiety levels and substance use, but since we showed within-subject changes in arousal in the same imaging session these are likely balanced within a run. Further work is needed to understand fluctuations in arousal over longer periods of time (e.g., days, months) and to relate these measurements to other quantifiable modalities (e.g., salivary cortisol measurement(Page et al., 2009)). It also may be interesting to explore if any specific arousal levels (and their associated hub disruptions at specific arousal levels help improve the performance of connectome-based fingerprinting and predictive modeling of individual traits or task performances. To compare arousal level-dependent brain network organizations between resting state and naturalistic paradigms may help to understand why naturalistic paradigms provides a better outcome in predicting behavior in some studies(Finn and Bandettini, 2021).

In conclusion, using the simultaneous measurements of resting state fMRI and pupillometry, we show evidence of a brain-wide decrease in between-network integration and a decrease in inter-subject variability of connector hubs at low relative to high arousal. Our results demonstrate that the estimation of *k*-hubness using SPARK, which reflects the number of overlapping networks for each node, is sensitive to the level of arousal within the resting state. By studying connector hubs of hierarchical brain network organizations, we suggest that modulations of arousal are not localized to specific brain areas, but rather have a more extensive, brain-wide impact that involves high-level communication between networks. Delineating arousal effects on functional connectivity reconfigurations may help advance future studies on the brain-behavior associations and neurological and psychiatric disorders where arousal may play a role in clinical phenotypes.

## 5. Acknowledgments

KL is supported by National Institute of Health (NIH) grants 5R01MH111424 and P50MH115716; CH is supported by a Medical Scientist Training Program training grant (NIH/NIGMS T32GM007205).

## 7. Data and Code Availability Statement

MRI preprocessing and visualization were performed using BioImage Suite version 3.5a1 and BioImage Suite Web (https://bioimagesuiteweb.github.io/webapp/index.html). The MATLAB scripts for SPARK implementation are available on https://github.com/multifunkim/spark-matlab. Several additional MATLAB scripts for SPARK adapted for this study are available on https://github.com/Kangjoo/arousal-rsfMRIpupil-hub. Imaging and pupillometry data are freely available on https://openneuro.org/datasets/ds003673/versions/1.0.0.

