## Supplementary materials for "Arousal impacts distributed hubs modulating the integration of brain functional connectivity"

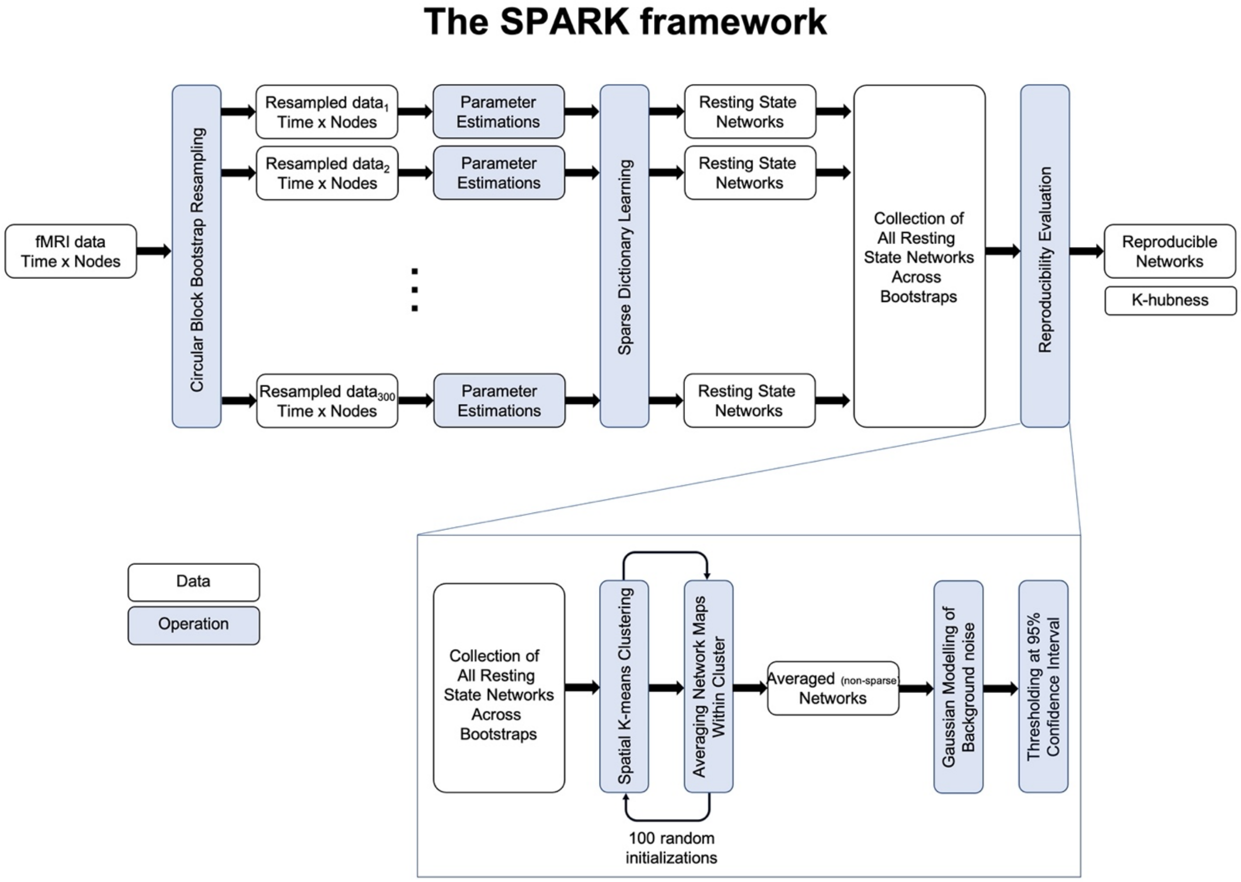


**Fig. S1.** A summary of SPARK method.

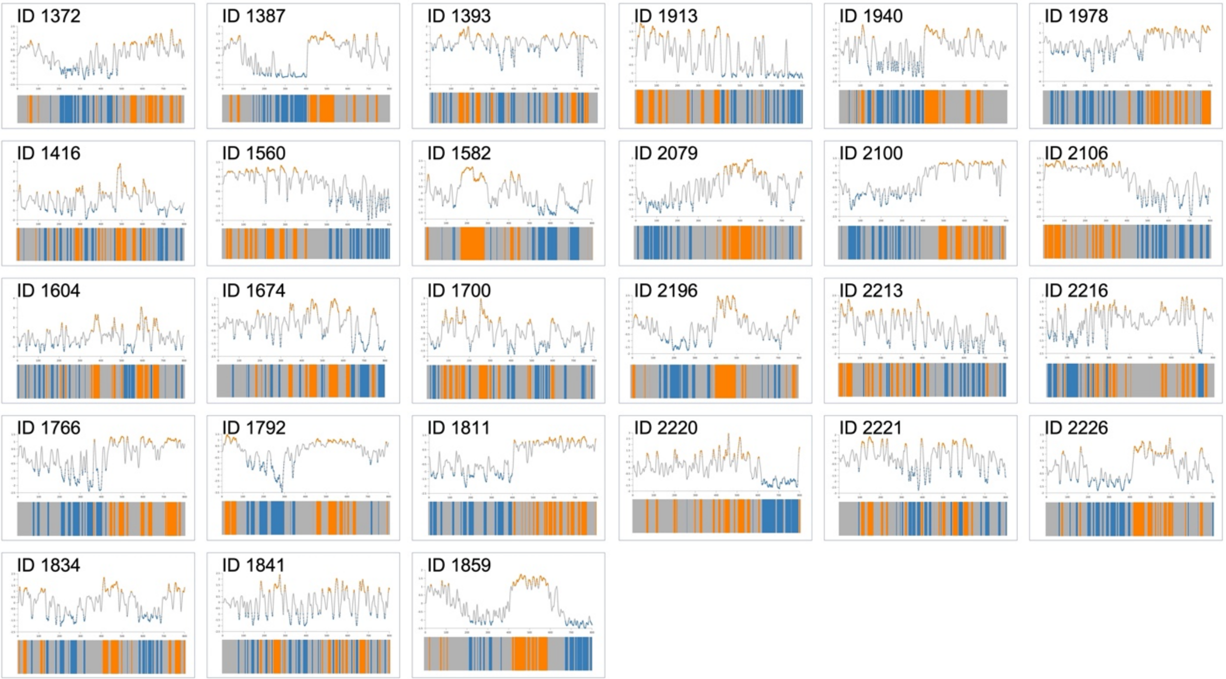


**Fig. S2.** Pupillometry time-courses and arousal state stratification (high arousal in orange and low arousal in blue) from individual subjects.

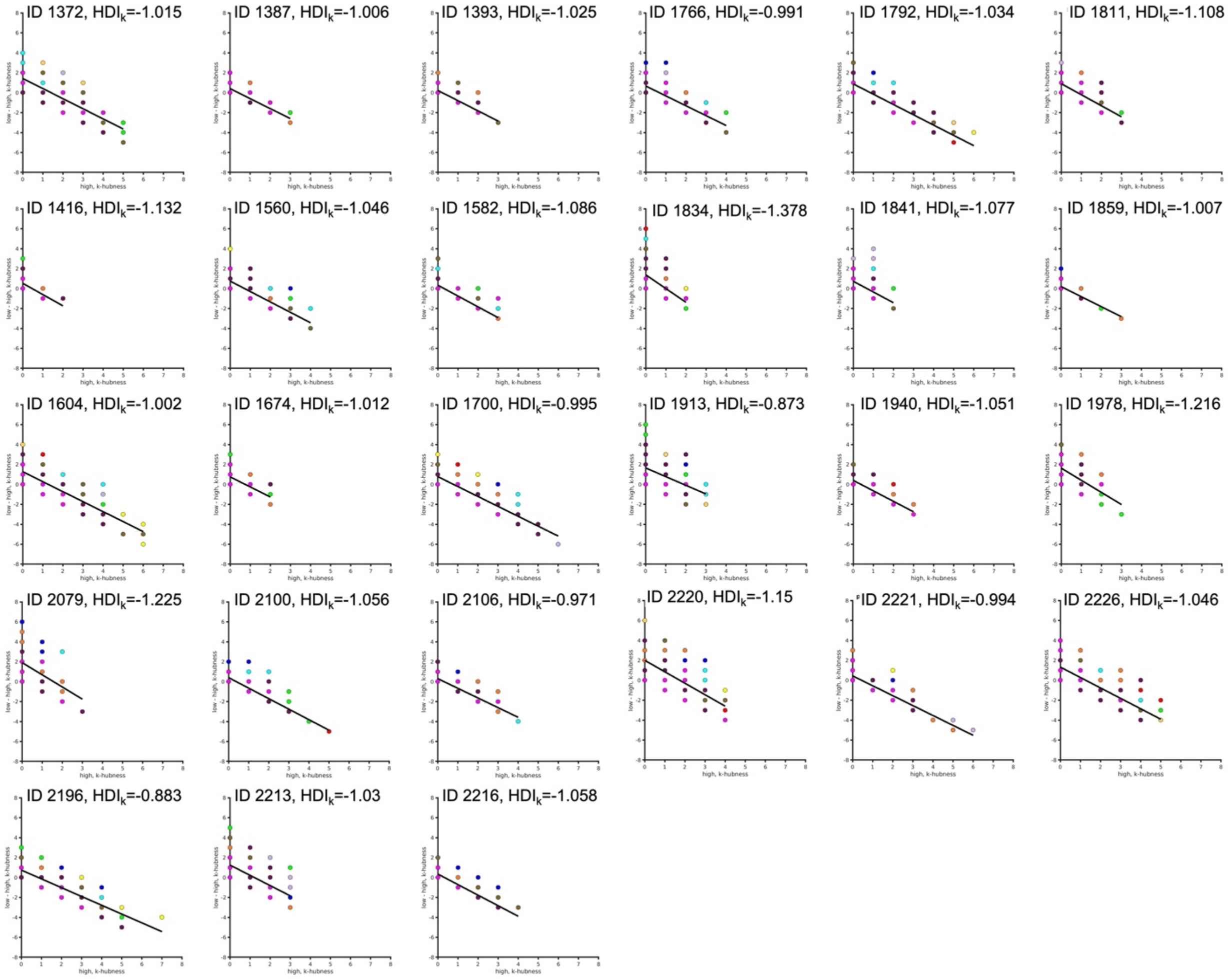


**Fig. S3.** Hub disruption index for k-hubness ($HDI_k$) estimated at the single subject level (N=27)

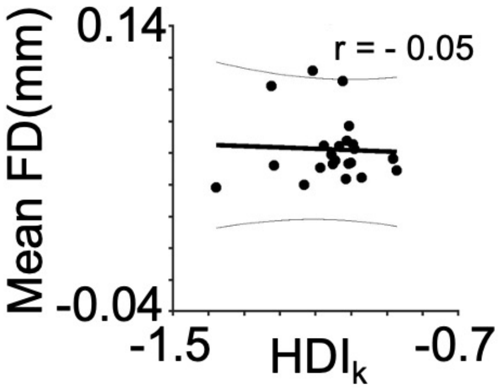


**Fig. S4.** No effect of head motion.

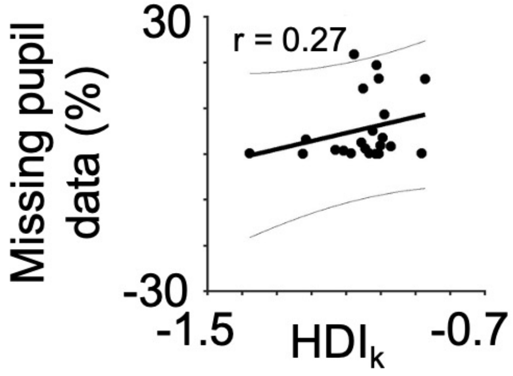


**Fig. S5.** No effect of eye-closure related loss of pupillometry data.

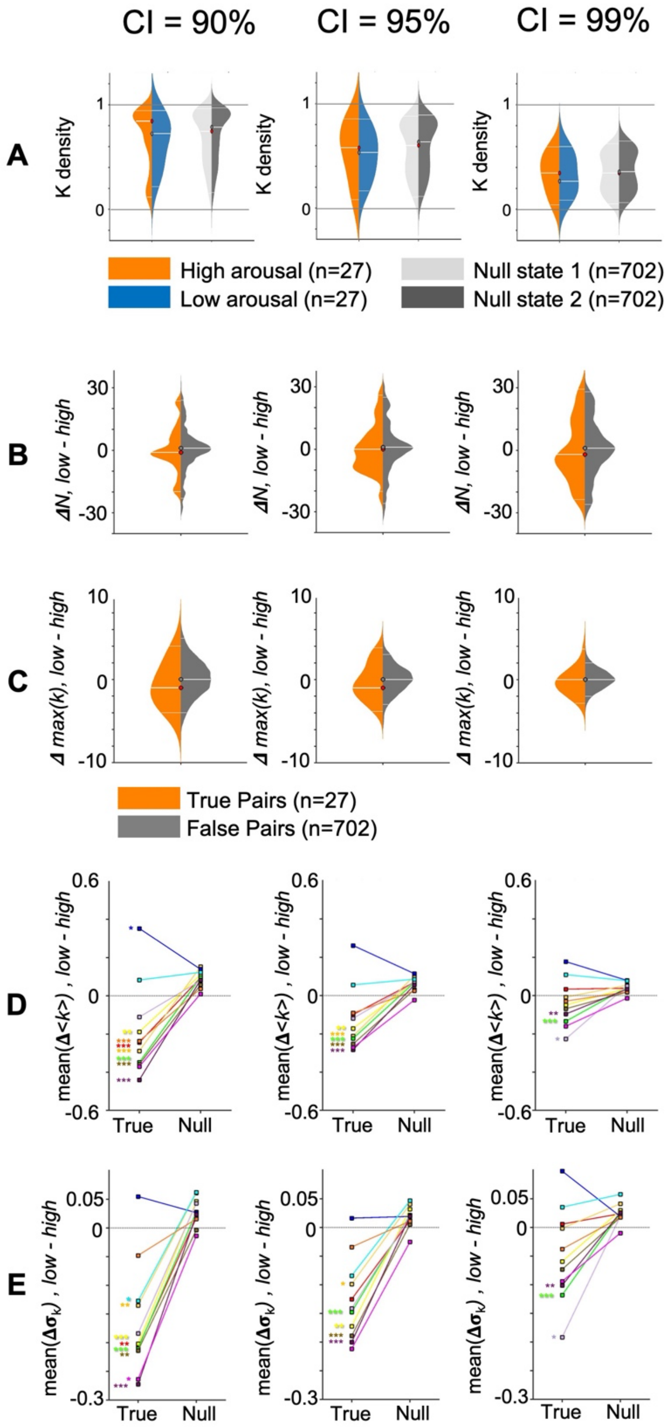


**Fig. S6.** Robustness of hub estimation using different thresholds (Confidence Interval, CI) for the reliability evaluation step within the SPARK framework.

**Table S1.** Demographics and data quality control (QC) metrics.

| Subject ID | Age | Sex | Handedness | Rest 1: EPI Mean frame-to-frame displacement (mm) | Rest 1: Eye-Tracking eye-closure artifacts (%) | Rest 2: EPI Mean frame-to-frame displacement (mm) | Rest 2: Eye-Tracking eye-closure artifacts (%) |
|---|---|---|---|---|---|---|---|
| **1372** | 31 | F | R | 0.044328 | 0.1123 | 0.042753 | 0 |
| **1387** | 35 | M | R | 0.081707 | 32.9378 | 0.072324 | 0 |
| **1393** | 29 | M | R | 0.090405 | 4.3777 | 0.120075 | 5.7365 |
| **1416** | 37 | F | R | 0.039752 | 0.8218 | 0.039885 | 0.9257 |
| **1560** | 21 | F | R | 0.06162 | 1.3187 | 0.049 | 1.0815 |
| **1582** | 22 | F | R | 0.057545 | 0.1975 | 0.04373 | 0 |
| **1604** | 27 | F | R | 0.04703 | 0.5285 | 0.060856 | 3.103 |
| **1674** | 22 | M | L | 0.056752 | 11.0008 | 0.078484 | 27.8292 |
| **1700** | 24 | F | R | 0.0604 | 3.3043 | 0.07019 | 3.7435 |
| **1766** | 22 | M | R | 0.059174 | 17.1137 | 0.066106 | 0.1123 |
| **1792** | 30 | M | R | 0.063826 | 0.24 | 0.064842 | 0 |
| **1811** | 29 | M | L | 0.092673 | 0.256 | 0.1311 | 1.0927 |
| **1834** | 24 | F | R | 0.038476 | 0 | 0.038218 | 0.30325 |
| **1841** | 27 | F | R | 0.059436 | 26.0775 | 0.069907 | 17.477 |
| **1859** | 27 | M | R | 0.031553 | 0.1425 | 0.074828 | 0 |
| **1913** | 24 | F | R | 0.034066 | 0.45375 | 0.063773 | 32.2682 |
| **1940** | 23 | M | R | 0.054733 | 28.4655 | 0.051066 | 0.15175 |
| **1978** | 22 | F | R | 0.045672 | 5.932 | 0.058694 | 0.4675 |
| **2079** | 24 | F | R | 0.106166 | 0 | 0.098454 | 0 |
| **2100** | 28 | F | R | 0.051148 | 2.769 | 0.066422 | 2.3247 |
| **2106** | 27 | M | R | 0.043171 | 0 | 0.045579 | 3.2692 |
| **2196** | 29 | F | R | 0.046251 | 0.28425 | 0.066404 | 0 |
| **2213** | 28 | M | R | 0.08281 | 5.1605 | 0.091175 | 8.97 |
| **2216** | 23 | F | R | 0.060108 | 11.7525 | 0.035896 | 4.9015 |
| **2220** | 31 | M | R | 0.0453 | 0.11225 | 0.042464 | 0 |
| **2221** | 25 | F | R | 0.045074 | 9.3082 | 0.052533 | 2.9732 |
| **2226** | 25 | F | R | 0.038957 | 0 | 0.038563 | 0.17525 |

**Table S2.** Nodes that showed decreases in k-hubness (node-wise bootstrap two-sample test, $p_{FDR}<.05$).

| Node Number (Shen et al., 2013) | Node definition (Shen et al., 2013) | Network definition (Noble et al., 2018) | Brodmann Area | MNI coordinate | Test statistic |
|---|---|---|---|---|---|
| 61 | R-Temporal | Motor | PrimAuditory (41) | (59.18, -2.26, 2.74) | -2.81 |
| 84 | R-Limbic | Motor | VentAntCing (24) | (5.25, -1.35,56) | -2.79 |
| 118 | R-Cerebellum | Cerebellum | Cerebellum | (33.77, -47.23,-53.93) | -2.47 |
| 154 | L-Prefrontal | Fronto-parietal | dlPFC(lat) (46) | (-42.97,42.04,11.04) | -2.56 |
| 162 | L-MotorStrip | Medial-Frontal | PreMot+SuppMot (6) | (-9.06,0.38,66.53) | -2.86 |
| 221 | L-Limbic | Limbic | VentAntCing (24) | (-5.1, 13.15, 28.74) | -2.86 |
| 229 | L-Limbic | Basal Ganglia | Hippocampus | (-21.49, -36.87,5.75) | -3.37 |
| 251 | L-Cerebellum | Limbic | Cerebellum | (-10.3, -37.72, -24.07) | -2.95 |
